# Multi-Parametric Profiling of Endothelial Cell Networks Reveals Functional Role of Glutamate Receptors in Angiogenesis

**DOI:** 10.1101/2019.12.25.888479

**Authors:** Heba Z. Sailem, Ayman Al Haj Zen

## Abstract

Angiogenesis plays a key role in several diseases including cancer, ischemic vascular disease, and Alzheimer’s disease. High throughput screening of endothelial tube formation provides a robust approach for identifying drugs that impact microvascular network formation and morphology. However, the analysis of resulting imaging datasets has been limited to a few phenotypic features such as the total tube length or the number of branching points. Here we developed a high content analysis framework for detailed quantification of various aspects of network morphology including network complexity, symmetry and topology. By applying our approach to a high content screen of 1,280 drugs, we found that many drugs that result in a similar phenotype share the same mechanism of action or common downstream signalling pathways. Our multiparametric analysis revealed a group of drugs, that target glutamate receptors, results in enhanced branching and network connectivity. Using an integrative meta-analysis approach, we validated the link between these receptors and angiogenesis. We further found that the expression of these genes correlates with the prognosis of Alzheimer’s patients. In conclusion, our work shows that detailed image analysis of complex endothelial phenotypes can reveal new insights into biological mechanisms modulating the morphogenesis of endothelial networks and identify potential therapeutics for angiogenesis-related diseases.

## Introduction

The primary function of microvascular networks across the body is to provide nutrients and oxygen supply for tissues. Both the vasculogenic and angiogenic processes work co-ordinately to form the network of tissue microvasculature. Vasculogenesis is the de novo formation of vessels from the assembly of progenitors or mature endothelial aggregates (Davis et al., 2011). While in angiogenesis, the new vessels are formed from pre-existing ones (Senger and Davis, 2011). Endothelial cell branching and tubulogenesis are critical elements for the formation of functional microvascular networks during embryonic development and postnatal life. Defects in vascular network structure or formation play a significant role in many pathological conditions, including ischemic vascular disease, cancer, neurodegenerative diseases, inflammatory disorders, and diabetes (Folkman, 2007). Consequently, the identification of small molecules or signalling components that regulate this complex process has important implications for many diseases.

The features of the nascent endothelial network are determined by genetic components and the surrounding microenvironment. Changes in these factors can affect vessel morphology, structure, organisation, or permeability leading to different pathologies (Folkman, 2007; Potente et al., 2011). For instance, the vessels that are formed in tumours are often abnormal and can have various phenotypes, including irregular organisation, loosely assembled vessel wall, abnormally wide or thin vessels, and tortuous serpentine-like morphology (Potente et al., 2011). On the other hand, leaky vessels have been implicated in diabetic retinopathy and neurodegenerative diseases (Folkman, 2007; Muzny et al., 2012). Developing methods for quantifying different aspects of endothelial networks is important for decoding the signalling components contributing to vascular morphology (Potente et al., 2011).

Endothelial cells have a remarkable ability to form tube networks, even in vitro cell cultures autonomously. The presence of vascular growth factors and appropriate extracellular matrix substrate induce a specific response in the endothelial cells to assemble and form branched tubes. For instance, within a few hours of being cultured on Matrigel, a natural basement membrane matrix, endothelial cells form branched networks (Kubota et al., 1988). This process mimics many facets of in vivo angiogenesis, including cell-cell interaction, elongation, motility, and cell-matrix adhesion. Therefore, the changes detected in endothelial network morphology reflect broadly molecular and cellular functions that propagates to specify tissue-level phenotypes.

High content imaging has proved to be a powerful approach to drug discovery. Target-based screening has been successful in identifying small molecules targeting a specific biological marker or process such as migration, proliferation, viability, or branching (Gallardo et al., 2015; Kalén et al., 2009; Wagner et al., 2018; Yarrow et al., 2005). However, it has been shown that monitoring morphological changes at the cellular level can confer predictive insights on the drug mechanism of actions and its effect on cell biology (Futamura et al., 2012). Detailed morphology analysis further allows detection of potential side or undesired drug effects, which ultimately can reduce the attrition rate of drug development pipelines and result in identifying more relevant therapeutic targets (Breinig et al., 2015). Furthermore, profiling of the heterogeneity of cellular responses upon exposure to drugs has been shown to be useful for further characterisation of the drug mechanism of action (Perlman et al., 2004; Slack et al., 2008). The screening of more complex cell culture models that mimic *in vivo* biology is starting to emerge as it is expected to select drugs that are more likely to mature in the clinic (Mills et al., 2019; Tung et al., 2011). Although vascular morphogenesis assays have been established (Al Haj Zen et al., 2016; Isherwood et al., 2013), a comprehensive analysis of these complex culture models are still lacking. Moreover, the extent that higher level tissue properties reflect those at the molecular level is yet to be established.

We propose that detailed phenotyping of endothelial network features in chemical genetic screening can aid the identification of drugs mechanism of actions and elucidate genetic components deriving this complex process. Specifically, we developed a quantitative multi-parametric method for monitoring complex phenotypic aspects of endothelial networks such as branching, connectivity, topology, and symmetry. This enabled in-depth angiogenic phenotype characterisation of drugs in the LOPAC_1280_ library that have a diverse mechanism of actions and impact most signalling pathways (Kalén et al., 2009). Importantly, through our multivariate analysis, we identified a novel antiangiogenic role for a group of glutamate receptors based on the network phenotype of their chemical inhibition by glutamate receptor antagonists. These results were validated using an orthogonal clinical dataset measuring gene expression in Alzhiemer’s Disease patients, which showed that these genes have lower expression in patients with worse prognosis. Therefore, these genes might provide a promising target in Alzheimer’s disease. The multi-parametric profiling of endothelial networks is not only relevant to better understand signalling components involved in the multistep endothelial network formation process but also hold the promise of discovering novel therapeutic targets.

## Results

### Identifying multi-dimensional phenotypic signatures of endothelial cell network formation

To comprehensively profile endothelial network patterns upon drug treatments, we used a dataset from a previous high content screen of LOPAC_1280_ small molecule library (Al Haj Zen et al., 2016). In the previous study, we identified 15 enhancers and 79 inhibitors of tube formation where the analysis have been limited to one feature of the network (total tube length). Despite the importance of this parameter as a descriptor of network formation, single parameter analysis does not account for other aspects of endothelial network formation. To comprehensively profile network morphology, we classified network elements as tubes, nodes (i.e., branching points), or meshes (area enclosed within tubes) (**Fig. 1A)** and (Carpentier et al., 2012). Tubes are further sub-classified into twigs, segments, and isolated tubes based on their connectivity while nodes are subclassified into junctions, master junctions, and extremities (**Methods**). In total, we extracted 106 features quantifying tube formation (e.g., number of tubes, and total tube length), tube morphology (mean tube length and thickness), mesh formation (number of meshes, mean mesh area, and total mesh area), and network connectivity based on the number of connected components and isolated tubes. Furthermore, we developed various global measures of the formed network: network symmetry (e.g., Standard Deviation (SD) of tube and mesh measurements), network complexity based on fractal analysis and Voronoi tessellation, and network topology based on network centrality measures from graph theory such as the length of the average length of shortest paths between nodes and minimum spanning tree (**Fig. 1A, Supplementary Table 1 and Methods**).

**Figure 1.**
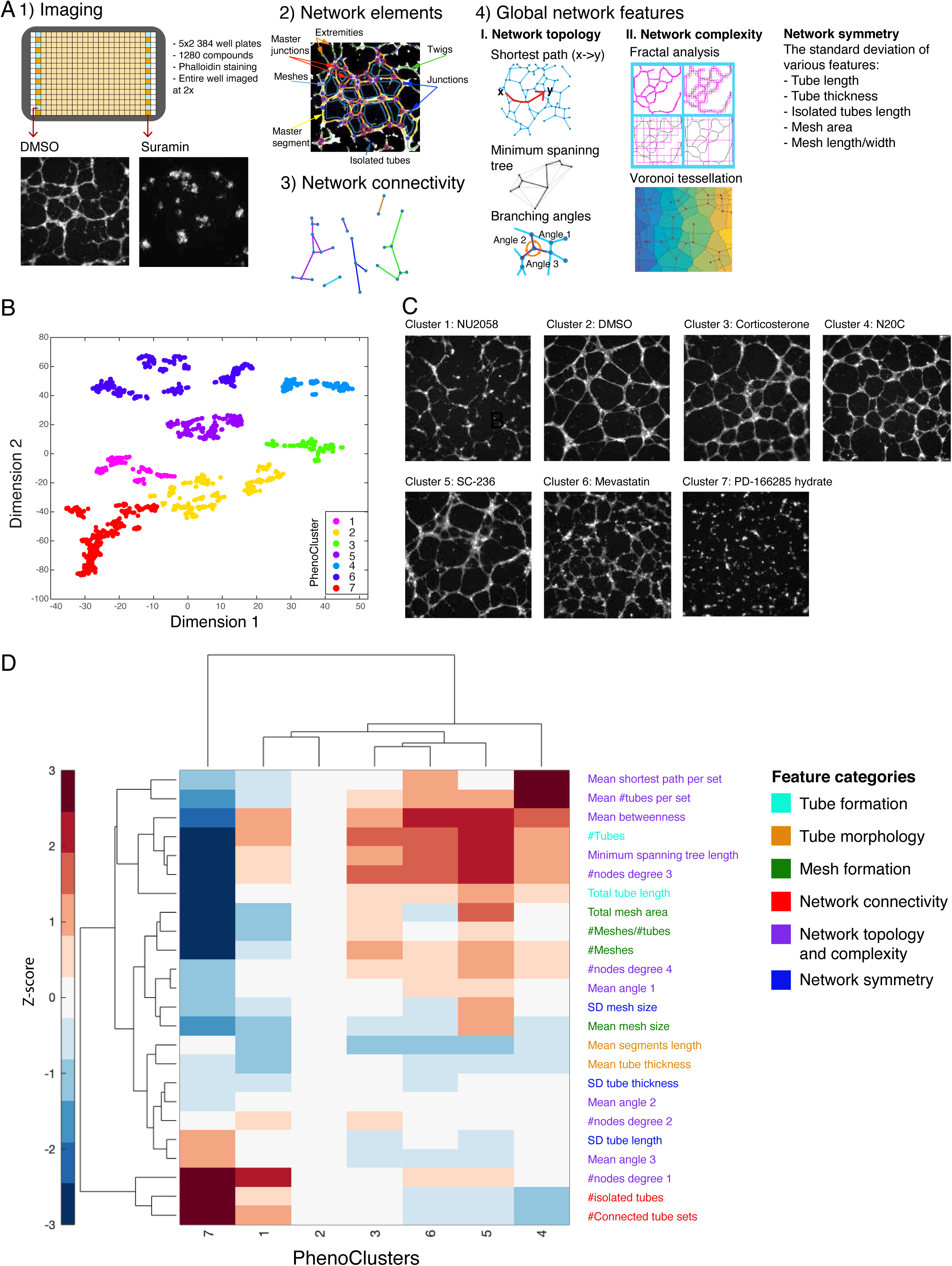
Quantitative analysis of vascular network phenotypes based on tube formation assay. (**A**) Image analysis pipeline involves network detection, classification of network elements and feature extraction. Cells are stained with Phalloidin and imaged at 2x. Representation of measured features including network topology, symmetry and complexity are indicated. **(B**) Clustering of LOPAC compounds in t-SNE space based on 106 phenotypic features. (**C**) Representative images for each PhenoCluster. (**D**) Quantitative differences between various clusters based on representative set of features from different categories.

Many drugs affected multiple aspects of endothelial network formation. Hierarchical clustering was performed to group small molecules with similar phenotypic profiles, using ward linkage and Euclidean distance (**Fig. 1B**). This analysis revealed seven distinct phenotypic clusters (PhenoClusters), where PhenoCluster 2 is composed of control-like drugs with no significant change in network features. Other clusters exhibited distinct phenotypes ranging from changes in branching and mesh formation to changes in network topology (**Fig. 1B-C and Supplementary Table 2**). Drugs in cluster 1 and 7 are characterised by significant disruption of features measuring network formation. However, only drugs in PhenoCluster 7 significantly decreased tube formation by endothelial cells (**Fig. 1C-D**). Many of the drugs in this cluster are potent inhibitors of angiogenesis such as the positive control Suramin and the VEGF inhibitors; DMH4, SU4312, SU5416 and Sunitinib (**Fig. 1D**). Moreover, 95% of the drugs that were identified as inhibitors in the previous study (Al Haj Zen et al., 2016) are in this cluster (75/79 Fisher exact test; *p*<1.45e-48).

Drugs in PhenoCluster 3, 4, 5, and 6 resulted in an enhancement of many tube and mesh formation features but varied in other features (**Fig. 1D, Sup Fig. 1A-B**). This increase in branching is also associated with improved connectivity in these clusters with except to PhenoCluster 3 that instead exhibited an increase in the number of 2-way junctions (**Supp. Fig. 1C, D, N**). PhenoCluster 4 is characterised by the highest increase in connectivity as most tubes belong to the same network which is reflected by the significant decrease in the number of connected sets and the number of isolated tubes (**Fig. 1D and Sup Fig. 1C-E**). On the other hand, drugs in PhenoCluster 5 affected mesh formation where the average and SD of mesh area are increased (**Fig. 1F-I**). They also affected the network topology as the orientation of tubes in 3-way nodes (i.e., branching points with three tubes) is changed where the largest angle is increased, and the smallest angle is decreased (**Sup Fig. 1J-K**). The network in PhenoCluster 6 showed a decrease in mesh area and increased in thin and short tubes which might indicate immaturity of these tubes (**Fig. 1C and Sup Fig. L, M, Q**). Taken together, these results illustrate that our multi-dimensional profiling allows distinguishing different morphologies of endothelial cell networks induced by drug treatment.

### Mapping endothelial network signatures to drug mechanism of actions and chemical structure

Our phenotypic clusters show significant enrichment for drugs with a common mechanism of actions (FDR *P*-value <0.05) (**Fig. 2A and Supplementary Table 3**). For example, PhenoCluster 7, which is characterised by the complete disruption of tube formation, is composed of many drugs that interfere with processes that are required for endothelial cell proliferation and migration. At the molecular level, these include drugs that disrupt cytoskeletal reorganisation and cellular adhesion such as tubulin and focal adhesion kinase inhibitors and anti-proliferative drugs such as DNA topoisomerase I and II inhibitors. While PhenoCluster 5 is enriched for a mechanism of actions associated with anti-inflammatory response including cyclooxygenase (COX) inhibitors and kappa agonists. These results confirm that our phenotypic features are biologically relevant and allow identifying drugs associated with the same mechanism of actions.

**Figure 2.**
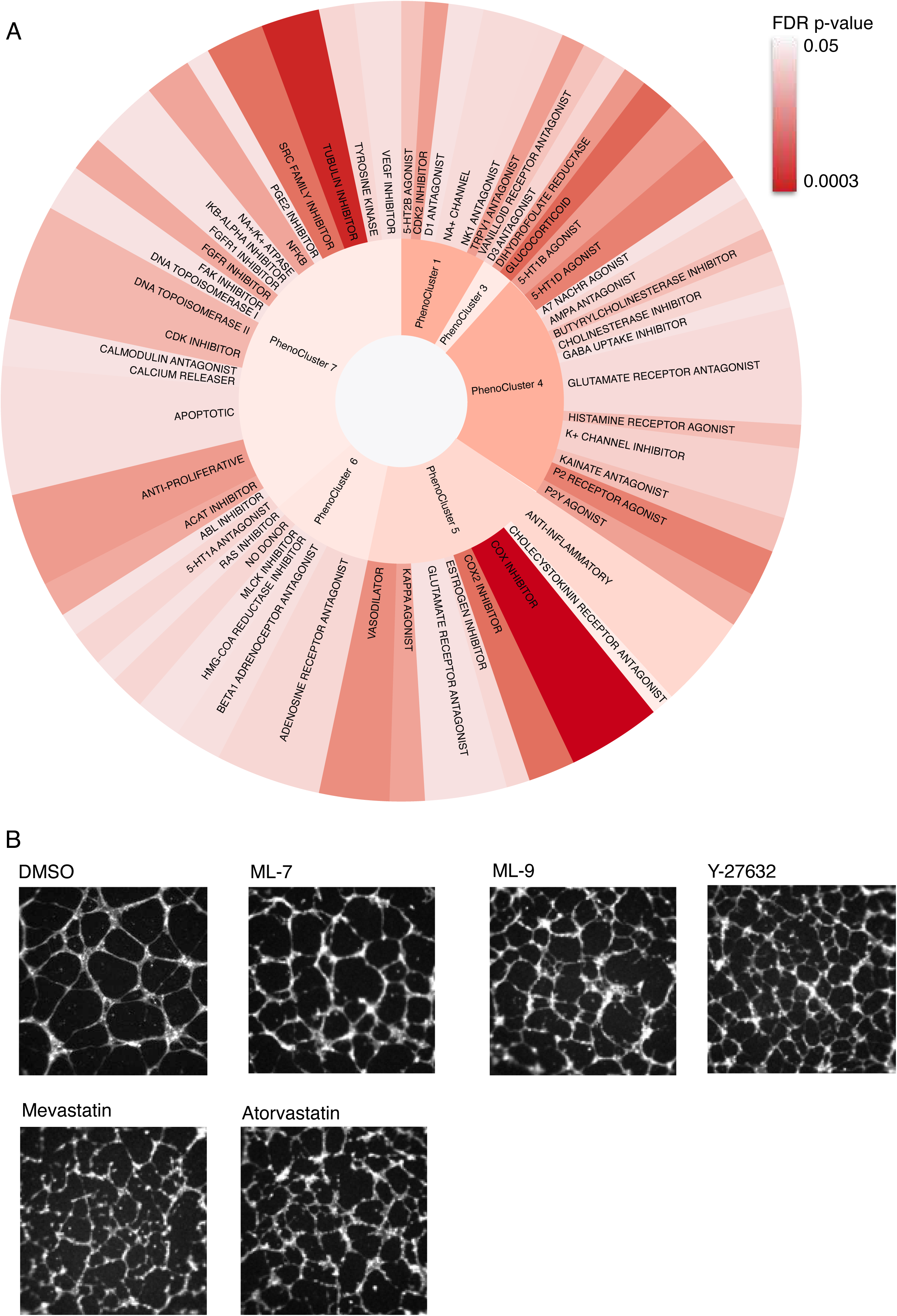
Cluster enrichment for mechanism of actions. (**A**) For each cluster, the enriched mechanism of actions are indicated proportional to their size. The significance of the enrichment is based on Fisher Exact test and corrected using false discovery rate. (**B**) Effects of drugs that inhibit Myosin Light Chain Kinase (ML-7 and ML-9), the ROCK inhibitor Y27632, and Statins (HMGA reductase inhibitors: Mevastatin and Atrovastatin).

Many compounds associated with Myosin Light Chain (MLC) signalling are in PhenoCluster 6, which is characterised by immature thin tubes. These include the Myosin Light Chain Kinase (MLCK) inhibitors MLC-7 and MLC-9, as well as the ROCK inhibitor Y-27632 (**Fig. 2A-B**). The observed phenotype is consistent with MLCK role in cell motility, polarization, and adhesion (Shen et al., 2010). Other myosin-related mechanisms that are enriched in this cluster are NO donor compounds (3 out of 4), Ras inhibitors (2 out of 2), and HMG-CoA reductase inhibitors (2 out of 2) (**Fig. 2B**). These results are in agreement with previous studies as NO production has been shown to inactivate MLCK (Kitazawa et al., 2009) and RAS has been shown to be required for MLCK activation (Nguyen et al., 1999). Interestingly, the inhibition of HMG-CoA reductase has been shown to reduce endothelial cell migration due to its effect on RhoA localisation, which is upstream of MLCK (Vincent et al., 2001). That drugs that target similar signalling pathways have a similar network signature illustrate that tissue level features can provide a robust proxy for changes at the molecular level. Next, we determined the chemical structural similarity between drugs in the same phenotypic cluster. Using Tanimoto distance (Breinig et al., 2015), we observe that some structurally similar drugs also have similar bioactivities and cluster together (**Fig. 3A-D, Fig. 3F, and Supplementary Table 4**). For example, the 5-HT2 serotonin receptor antagonist Ritanserin and Pirenperone in PhenoCluster 1 are structurally similar (**Fig. 3A**). Additionally, four adrenoceptor agonists in PhenoCluster 6 shared similar chemical structures, including Isoproterenol hydrochloride, Isoetharine mesylate, and Epinephrine hydrochloride (**Fig. 3B**). On the other hand, L-DOPS and (-)-alpha-Methylnorepinephrine are other adrenoreceptor agonists in PhenoCluster 6 that are structurally distinct from the aforementioned adrenoreceptors but rather are structurally similar to the Tyrosine hydroxylase inhibitor 3-Iodo-L-tyrosine (**Fig. 3C**). Therefore, our phenotypic clusters could, in some cases, be explained by structural similarity, bioactivity, or both, which highlight the importance of phenotypic profiling to identify drugs with similar effects on cell physiology.

**Figure 3.**
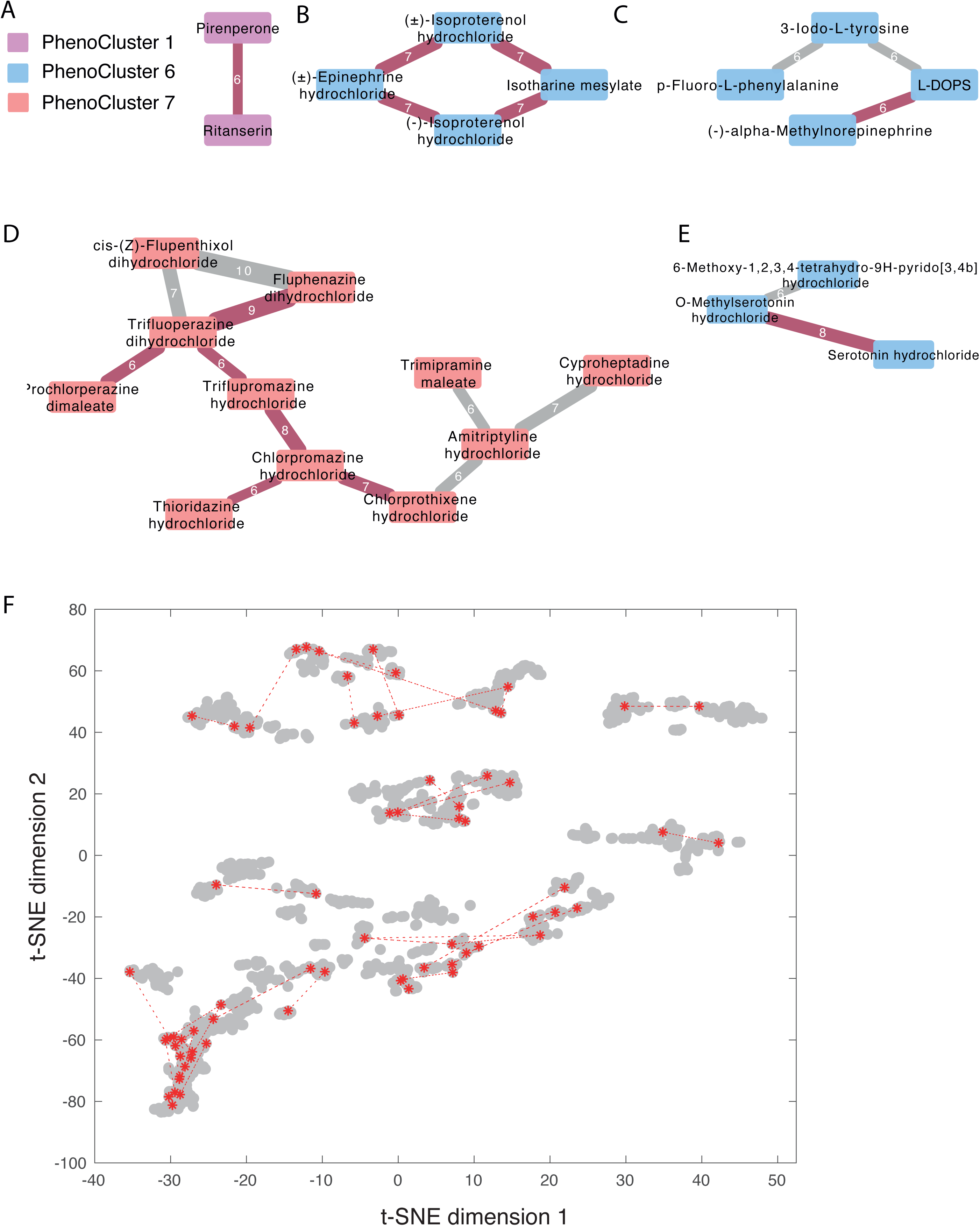
Compounds structural similarity. (**A-E**) structural similarity between compounds in the same cluster where drugs are represented as nodes and similarity as edges. Compounds are connected by a red edge if they share the same mechanism of action and a grey edge otherwise. The numbers on the edges indicate the z-scored structural similarities rounded to the nearest integer. Node colour indicates the compound cluster. **(F)** Compounds with a similar structure in the same PhenoCluster are connected by a dashed line.

Analysing structural similarity along phenotypic similarity allows identification of potential off-target effects or the mechanism of action of tested compounds. In particular, when a drug that is structurally similar to a group of drugs with a known mechanism of action, have also a similar phenotypic profile (Westfall et al., 1997). For example, we found a group of structurally similar drugs in PhenoCluster 7, many of which are antagonists of dopamine, including Triflupromazine hydrochloride, Chlorpromazine hydrochloride, Prochlorperazine dimaleate, Fluphenazine dihydrochloride (**Fig. 2D**). However, the serotonin receptor antagonist Cyproheptadine Hydrochloride, serotonin inhibitor Trimipramine maleate, and adrenoceptor antagonist Amitriptyline hydrochloride are structurally similar to the dopamine antagonists in this cluster. This might suggest an off-target effect for these drugs, which is consistent with the fact that these drugs can have unselective activity against dopamine (Young et al., 2005).

On the other hand, 6-Methoxy-1,2,3,4-tetrahydro-9H-pyrido[3,4b] indole is a monoamine oxidase inhibitor (MOAI) that is structurally and phenotypically similar to two serotonin receptor agonists (**Fig. 3E**). The phenotypic similarity between these drugs is consistent with the fact that MOAI inhibits serotonin metabolism resulting in its accumulation and activation of serotonin receptors in what is known as serotonin syndrome (Boyer and Shannon, 2005; Entzeroth and Ratty, 2017). Therefore, combining phenotypic and structural similarity can facilitate the determination of the likely affected mechanism of action of a drug in a specific biological context.

### Glutamate receptor antagonists result in distinct multivariate endothelial network signatures

A group of drugs in PhenoCluster 4 is significantly enriched for several mechanisms of actions associated with glutamate neuroreceptors (**FDR p-value < 0**.**05, Fig. 2A**). This cluster showed mainly increased branching and connectivity (**Fig. 1D**). In particular, ten glutamate receptor antagonists out of 42 are in this cluster, three of which target both AMPA receptors and Kainate receptors (FDR p-value <0.05). A smaller group of glutamate receptor antagonists phenocopied PhenoCluster 5 (7 out of 42 compounds). Although drugs in this cluster resulted in increased branching, they also altered the network topology and mesh formation (**Fig. 1D**). Notably, the glutamate receptor antagonist memantine, which is used for treating Alzheimer’s patients, is in PhenoCluster 5. These results suggest that changes in glutamate receptor activity could affect the formation of microvasculature network.

Many mechanisms of actions related to glutamate neurotransmission were also enriched in PhenoCluster 4. For instance, inhibiting the uptake of GABA, which is a product of glutamate in the brain (Bak et al., 2006), results in a similar phenotypic signature. PhenoCluster 4 is also enriched for ATP-activated purinergic P2Y receptor agonists (3 out of 4 drugs and **Fig. 2A**), which has been shown to reduce the expression of AMPA receptors (Zonouzi et al., 2011). Furthermore, the serotonin agonists of both 5-HTR1B and 5-HTR1D receptors, but not 5HTR2B, which is in PhenoCluster 1, are also enriched in this cluster (p-value < 0.001). This in line with the fact that 5-HTR1 inhibits glutamate and GABA release, while 5-HTR2 has the opposite effect (Ciranna, 2006). Interestingly, P2Y agonists and HTR1B serotonin receptors have been shown to play a proangiogenic role (Cooke and Ghebremariam, 2008; Gerasimovskaya et al., 2008; Iwabayashi et al., 2012). Altogether these results further support that glutamate receptors and related neuroreceptors play a role in angiogenesis.

### Integrative meta-analysis reveals a link between glutamate receptors and angiogenesis in patients with Alzheimer’s disease

Since glutamate receptors and angiogenesis are both implicated in Alzheimer’s disease, we sought to validate our finding that glutamate receptors activation state affects angiogenesis. We utilised a microarray dataset measuring gene expression in 301 Alzheimer’s patients (Bennett et al., 2018; Willard and Koochekpour, 2013). The dataset is composed of brain samples with different Braak stages (1-6), where stage 6 is the most severe form of the disease (Braak and Braak, 1995). The dataset spans four Brodmann (BM) brain regions: Frontal Pole (BM10), Superior Temporal Gyrus (BM22), Parahippocampal Gyrus (BM36), and Inferior Frontal Gyrus (BM44). To define an angiogenic signature, we investigated a set of known pro-angiogenic genes that were previously described to be involved in cancer and Alzheimer’s pathogenesis (Bennett et al., 2018; Masiero et al., 2013).

First, we determined whether the selected set of pro-angiogenic genes are associated with patient outcomes. We computed the fold change of expression values of these genes in high Braak stage (5 and 6) versus low Braak stage (1 and 2) in different brain regions (**Fig. 4A and Supplementary Fig. 2A**). The classically affected Superior Temporal Gyrus (BM22) and Parahippocampal Gyrus (BM36) regions show the highest fold-change in the expression of many pro-angiogenic genes. This is in accordance with a previous study (Bennett et al., 2018). TIE1, ITGB5, ESAM, S1PR1, CDH5, VWF, KDR are among the genes with the highest expression fold-change in patients with high Braak scores (Fold change >1.2, Kolmogorov–Smirnov p-value < 0.001). These results confirm a link between the expression of pro-angiogenic genes and the prognosis of Alzheimer’s patients.

**Figure 4.**
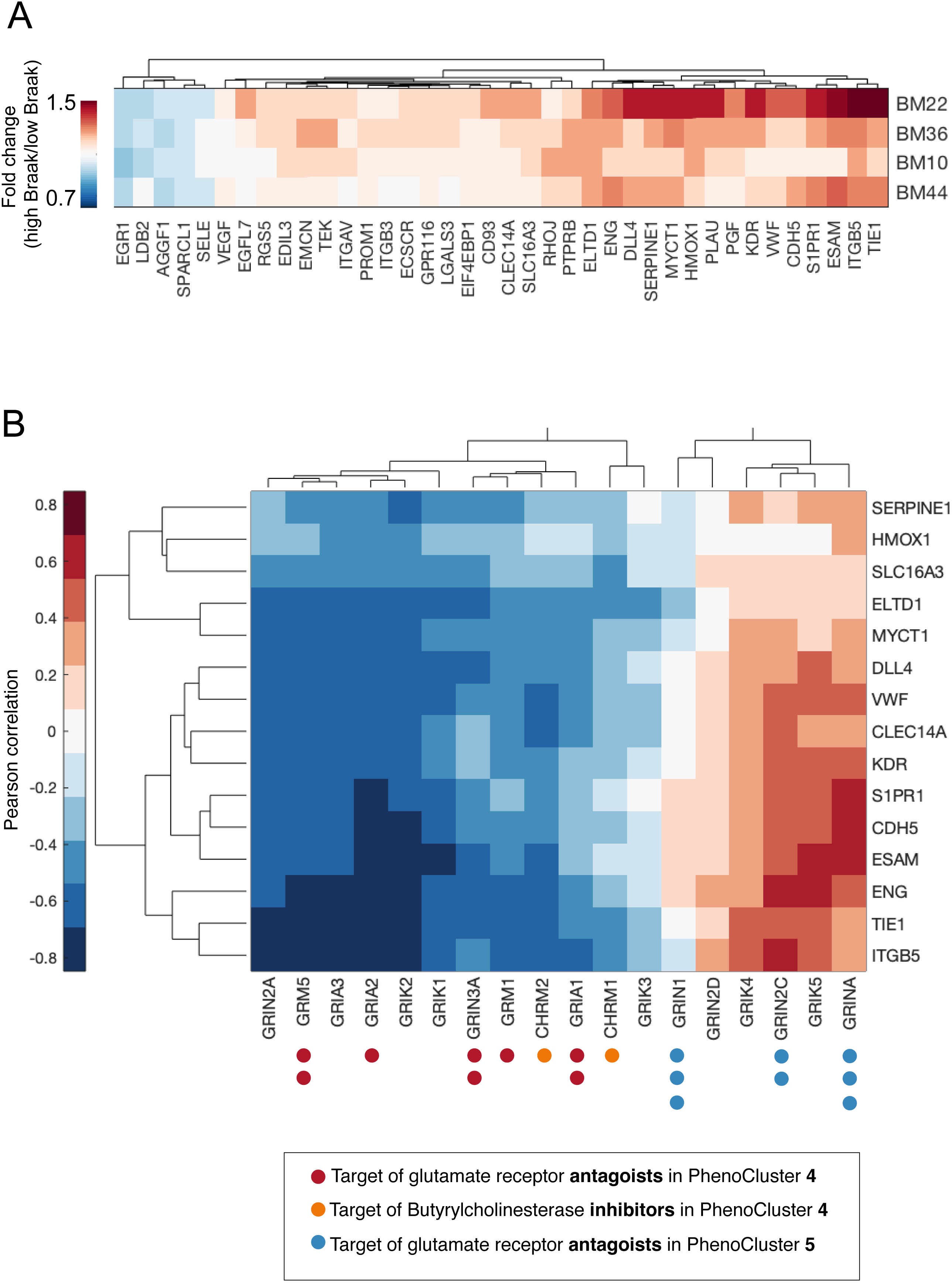
Association between glutamate receptors expression and angiogenesis in patients with Alzheimer’s disease. (**A**) Fold change of pro-angiogenic gene expression in high Braak stage Alzheimer’s patient. (**B**) Pearson correlation between the expression of various glutamate receptors and pro-angiogenic genes that have higher expression in Braak stage 5 or more. The number of dots next to the genes represent the number of drugs that target these genes. Dot colour indicates mechanism of action and PhenoCluster. Target genes of butyrylcholinesterase inhibitors that exhibited PhenoCluster 4-like phenotype were included for comparison.

Next, we determined the correlation between angiogenesis and glutamate receptor genes in Alzheimer’s patients. In this analysis, we included target genes of glutamate receptors antagonists in PhenoCluster 4, such as GRM5 and GRIN3A, as their chemical inhibition shows a “pro-angiogenic” phenotype (**Methods** and **Supplementary Table 5**). We also included target genes of butyrylcholinesterase inhibitors that induced PhenoCluster 4-like phenotype for comparison (**Fig. 4B; Supplementary Table 5**). We focused our analysis on Superior Temporal Gyrus (BM22) as it showed the greatest change in the expression of pro-angiogenic genes. We observed that the expression of many glutamate receptors in PhenoCluster 4 is negatively correlated with pro-angiogenic genes (Pearson coefficient<-0.3, and p-value<1.5e-05). Likewise, the expression of CHRM1 and CHRM2 genes, which are inhibited by the butyrylcholinesterase inhibitor Ethopropazine hydrochloride, is also correlated negatively with the expression of pro-angiogenic genes (p-value < 0.05). These results show that chemical genetic perturbations of genes that result in a similar network phenotype also have similar transcriptional profiles *in vivo*, which further confirm the validity of our high content analysis. Moreover, these results further support an anti-angiogenic role for a group of glutamate receptors, including GRM5 and GRIN3A.

On the contrary, the glutamate receptors GRIN1 and GRINA that are antagonized by drugs in PhenoCluster 5 correlated positively with pro-angiogenic genes (**Fig. 4B** and **Supplementary Table 5**). Similar correlation patterns are observed in the other brain regions except for Inferior Frontal Gyrus (BM44) (**Sup Fig. 2B-D**). These results support a differential role of glutamate receptors in angiogenesis, which can have an important implication for Alzheimer’s disease.

In order to evaluate the link between glutamate receptors expression and patient outcomes, we performed hierarchical clustering of Alzheimer’s patients based on the expression profiles of glutamate receptor genes. We identified three main patient clusters: P1-P3 (**Fig. 5A**). Cluster P1 is enriched for transcription profiles of the Inferior Frontal Gyrus region (65.38% of BM44 profiles) (**Fig. 5A-B**). Most glutamate receptors have moderate to high expression in this region. On the other hand, the expression of anti-angiogenic glutamate receptors identified by our analysis, have high expression in Cluster P2 (**Fig. 5A**). This cluster is almost void of profiles from BM44 region (**Fig. 5C**). In contrast, Cluster P3 exhibits a low expression of anti-angiogenic glutamate receptors (**Fig. 5A**). Interestingly, only Cluster P3 show significant enrichment for patients with high Braak stage where 59.44% of the patients in this cluster are stage 5 or 6 (**Fig. 5D**, Fisher’s exact test p-value < 2.2e-09 and Table 1). This enrichment is most significant in BM22 and BM36 brain regions (**Fig. 5E**, Fisher’s, and Table 1). Visualization of the expression profiles in Figure 5A in reduced dimensional space using t-SNE, shows that Cluster P1 and Cluster P2 are well separated while Cluster P3 deviate from these clusters. This might suggest a transition in the transcriptional state of glutamate receptors in Alzheimer’s patients, which correlate with a worse patient outcome (**Fig. 5G**). Taken together, our findings suggest that the down-regulation of glutamate receptors that are chemically inhibited in PhenoCluster 4 can contribute to Alzheimer’s pathogenesis and prognosis through increased angiogenesis.

**Table 1:** Enrichment of high Braak stage patients (grade 5 and 6) per patient cluster/region based on Fisher Exact Test

| Region | Cluster P1 | Cluster P2 | Cluster P3 |
| --- | --- | --- | --- |
| BM10 | 0.2207 | 0.9988 | 0.0075 |
| BM22 | 0.8736 | 0.9969 | 0.0007 |
| BM36 | 0.1611 | 1.0000 | 0.0007 |
| BM44 | 0.9709 | 0.1221 | 0.1137 |

**Figure 5.**
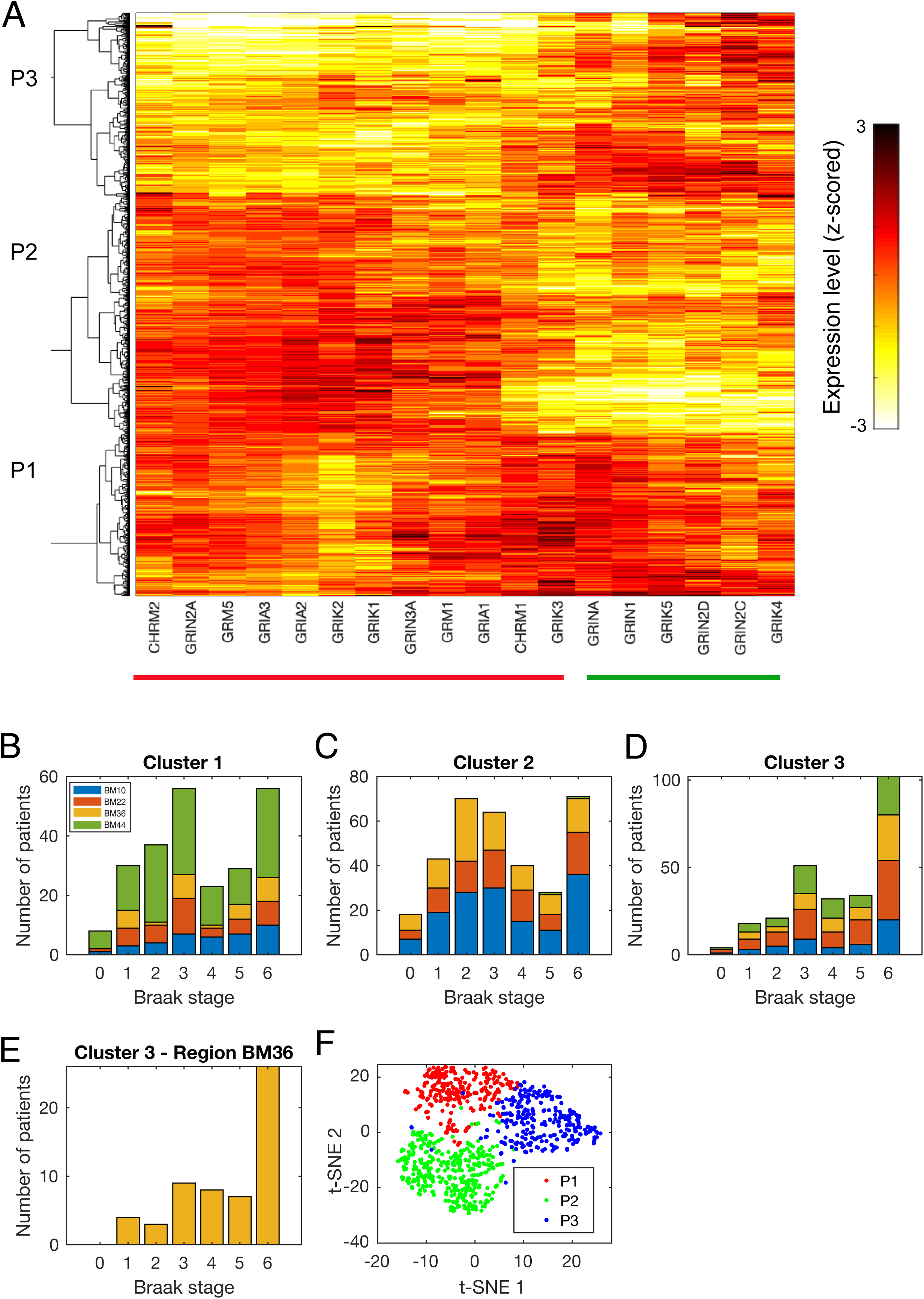
Analysis of glutamate receptors expression in Alzheimer’s patients. (**A**) Clustering of transcriptional profiles of glutamate receptors in different brain regions in Alzheimer’s patients shows three patient clusters. (**B-D**) Number of patients with different Braak stage in the different clusters. (**E-F**) Same as B and D but for brain region BM36. (**F**) Visualisation of transcriptional profiles in (A) using t-SNE.

## Discussion

The multicellular organisation of endothelial cells requires tight coordination of multiple molecular and cellular functions to form a microvascular network. Dysfunctional signalling pathways of endothelial cells at the molecular level should manifest in changes in network formation and morphology at the tissue level. Indeed, we illustrate that extensive phenotyping of endothelial networks upon chemical genetic perturbations can provide critical insights on mechanisms of actions and signalling components affecting the dynamics of self-organising endothelial cells. To our knowledge, our work provides the most comprehensive quantitative analysis of endothelial network phenotypes based on the largest screening dataset employing tube formation assay.

Our work demonstrates that quantitative phenotypic profiling of endothelial networks allows biologically relevant classification of cell responses to pharmacological perturbations. For example, compounds in PhenoCluster 6 displayed a network with small and thin tubes, which potentially could be due to the cell’s inability to form mature tubes. In this cluster, drugs that inhibit the ROCK and Myosin Light Chain Kinase (MLCK) have similar network phenotypes. Both ROCK and MLCK have a common downstream mechanism and reduce phosphorylation of myosin light chains (MLC). Myosin signalling is involved in many biological processes such as cell shape, motility, and cell-cell adhesion, that are critical for tube formation (Fischer et al., 2009). Interestingly, statins, which are inhibitors of liver HMG-CoA reductase, were phenotypically clustered with ROCK and MLCK inhibitors. Statins are FDA-approved drugs for lowering blood cholesterol (Zeng et al., 2005). However, it is becoming increasingly apparent that Statins have pleiotropic effects independent of cholesterol-lowering. Statins prevent the synthesis of isoprenoid intermediates that are necessary lipid attachments for the post-translational modification of small GTP-binding proteins, including RhoA, the primary activator of ROCK (Shen et al., 2010). Therefore, the phenotypic similarity of endothelial network morphology can be used for predicting non-canonical molecular mechanisms targeted by drugs in a particular biological context.

CDK2 inhibitors were enriched in PhenoCluster 1, which is characterised by poor network formation (Fig. 2A). This cluster can reflect the reduction of cell motility rather than cell proliferation. It has been shown that tube formation assay is proliferation-independent (Kubota et al., 1988). In our screen, the cells were fixed after 8 hours of drug treatment to capture direct compound effects. Therefore, we assume that the detected effects on the morphology of endothelial network is not affected by the potential change of endothelial cell number during the assay. Supporting our findings, recent studies reported that CDK2 could modulate cell migration (Andrés, 2004).

Imaging of cells at single-cell resolution can provide additional information on cell biology, however, it can be not feasible in large screening studies. Here we show that multivariate analysis of endothelial network phenotypes – without detailed single-cell quantification - can be sufficient for cellular profiling responses to drugs.

Multivariate analysis of network phenotypic features also revealed that glutamate receptor antagonists are significantly enriched in PhenoCluster 4, which resulted in enhanced network connectivity. This is an interesting finding since the glutamate is one of the most prevalent neurotransmitters in the central nervous system, but the link between glutamate receptors and angiogenesis remains unclear. It has been previously shown that endothelial cells express multiple glutamate receptor types, including NMDA, AMPA, and metabotropic biochemical receptors. Additionally, they are functional upon the exposure of glutamate excess (András et al., 2007; Krizbai et al., 1998; Sharp et al., 2003). Antagonists of both ionotropic and metabotropic receptors receptor types were found to results in similar phenotypic features of the endothelial network. Ionotropic glutamate receptors are ligand-gated ion channels that open to allow ions such as Na^+^, K^+^, Mg^++^, or Ca^++^ to pass through the membrane in response to glutamate binding (Traynelis et al., 2010). Interestingly, in our study, potassium channel blockers were also enriched in PhenoCluster 4. Modulation of potassium channel activity has been shown to influence endothelial cell functions, including angiogenesis (Brähler et al., 2009; Umaru et al., 2015). Thus, we speculate that the in vitro angiogenic effect of glutamate receptor inhibition is mediated by the changes of potassium transit via the glutamate-activated potassium channels.

The anti-angiogenic effect of some glutamate receptors might initially seem contradictory to the fact that the glutamate receptor antagonist memantine is used to normalise glutamate neurotransmission in Alzheimer’s patients (Zhang et al., 2016). However, our multi-parametric analysis shows that memantine phenocopied compounds in drug cluster 5. We found that GRIN1 and GRINA that are antagonised by multiple drugs in PhenoCluster 5, positively correlate with pro-angiogenic genes. Furthermore, GRINA and GRIN1 expression in patient cluster P3 is associated with worse patient outcomes, which is the opposite trend to glutamate genes that are targeted by drugs in PhenoCluster 4. These results suggest that GRIN1 and GRINA are necessary for intact network formation. However, more experiments are needed to validate the role and the effect of glutamate receptor antagonists in PhenoCluster 5, such as memantine, on single-cell function, and how the change in network topology *in vitro* corresponds to vascular morphology *in vivo*.

Our integrative analysis of glutamate receptor genes that are identified by our phenotypic screen shows that expression patterns of glutamate receptors in Alzheimer’s patients fall into two groups. In particular, expression of genes that are known to be antagonised by drugs in PhenoCluster 4 negatively correlates with pro-angiogenic genes, and their suppression is associated with worse patient outcomes. One limitation of our meta-analysis is that we cannot determine whether glutamate receptor genes identified through our phenotypic screen are expressed by endothelial cells based on bulk gene expression. Nonetheless, the striking similarity of glutamate receptors profiles based on their network morphology *in vitro* and their transcriptional profiles in Alzheimer’s patients strongly supports a differential role of glutamate receptors in angiogenesis. Moreover, our results can have important implications when targeting the glutamatergic system in Alzheimer’s patients.

During vascular development, endothelial and neuronal cells have been shown to share multiple regulatory proteins that guide the motility and growth of both neuronal and endothelial cells in development (Klagsbrun and Eichmann, 2005). Furthermore, neurotransmitters, such as glutamate, and serotonin have also been shown to play a role in neuronal guidance during vascular development (Ruediger and Bolz, 2007). In adulthood, many clinical studies reported the role of neurotransmitters such as serotonin and dopamine in tumour angiogenesis (Asada et al., 2009; Peters et al., 2014). Given the strong anatomical and functional relationship between endothelial cells and neural cells such as astrocytes and glial cells in the Blood-Brain Barrier (BBB), glutamate could play a role in endothelial cell biology where BBB is dysfunctional in diseases such as vascular dementia or Alzheimer’s disease (Montagne et al., 2015; Zlokovic, 2011). Our findings highlight a potential reciprocal relationship between dysregulated neurotransmission and angiogenesis in the context of Alzheimer’s disease. Further experimental studies are warranted to elucidate the molecular mechanisms underlying this relationship.

In conclusion, we show that accounting for different aspects of vascular network morphology is important to understand the biological mechanisms underlying the behaviour of complex biological systems. As endothelial cells are interweaved in almost all tissues, our work provides a valuable resource for drug discovery studies aiming at targeting angiogenic-related processes. We envision that our multiparametric phenotyping can increase the efficacy and sensitivity of angiogenic drug development and advance our understanding of endothelial cell biology.

## Methods

### Dataset

The dataset used in this study was generated by an image-based screen of 1280 compounds based on LOPAC library as described previously (Al Haj Zen et al., 2016). Briefly, HUVEC endothelial cells were cultured on top of Matrigel after adding compounds and incubated at 37°C, 5% CO_2_ for 8 hours. Then cells were fixed and stained with Phalloidin and imaged by the high content imaging system Operetta from Perkin Elmer.

### Image analysis

Image analysis was performed using ImageJ and MatLab.

#### Classification of network elements

Skeletonization was used to detect the network structure. Network skeleton is used to extract tubes and branching points (nodes) which are classified into 1) segments: tubes that are connected to the rest of the network from both sides, 2) twigs: tubes that are linked to the rest of the network from one side and 3) isolated tubes: tubes that are not connected to the rest of network, and 4) master segments: segments that are connected to other segments from both sides. Similarly, nodes are also subclassified into 1) junctions: nodes linking two or more tubes, 2) extremities: nodes that are linked to only one tube and 3) master junction: a junction that links at least three master segments. For each of these elements various statistics were computed including mean, standard deviation, number and total of each element length or area.

#### Network topology characterisation

Measurements from graph theory were used to quantify vascular network topology. The vascular network was represented as a graph where nodes in the endothelial network correspond to a set of vertices and tubes to a set of edges in the graph. Different centrality metrics of the graph were computed.

Voronoi tessellation was defined based on the branching points. Given a set of points in a spatial plane then the Voronoi diagram is the partitioning of a plane with a set of points into convex polygons such that each polygon contains exactly one generating point and every point in a given polygon is closer to its generating point than to any other. The average and standard deviation of the resulting polygons sizes can reflect the homogeneity of nodes distribution.

Texture measures based on Grey Level Co-occurrence Matrix (GLCM) were used to quantify intensity variation in Phalloidin.

### Clustering of phenotypic data

Empty wells and wells with artifact were identified manually and filtered from further analysis. All data were normalised to plate control by subtracting the mean of DMSO-profiles and dividing by the standard deviation of DMSO-profiles for that plate. Then compound replicates were averaged. All the resulting data were scaled between 0 and 1. Then t-SNE was used to reduce the dimensionality of the data to two dimensions. Drugs are then clustered using hierarchical clustering based on ward linkage and Euclidean distance.

### Drug library annotation

The drug information was downloaded from the DrugBank Database in June 2017. Each drug was annotated with multiple mechanisms of actions where it has multiple targets. Additionally, as drugs varied in the annotated mechanism of action specificity, drugs were annotated with the more general as well as specific mechanism of action. For example, drugs that are annotated as HTR1A receptors are also annotated with the more general mechanism of action HTR receptors.

### Glutamate receptors target identification

Target genes of glutamate receptor antagonists in PhenoCluster 4 and 5 were identified based on DrugBank database. Additionally, we used Sigma website which listed many potential targets. To increase the confidence of the association between gene-drug phenotype, only genes that are targeted by at least two drugs in the same PhenoCluster based on Sigma were indicated in Fig. 4B (Supplementary Table 5).

### Analysis of Alzheimer’s expression data

We used the publicly available Mount Sinai Medical Center Brain Bank (MSBB) gene expression dataset of 301 Alzheimer’s patients (Wang et al., 2018). The dataset is composed of brain samples with different Braak stages (1-6) that are based on the topographical distribution of neurofibrillary tangles and neuropil threads where stage 6 is the most severe form of the disease (Braak and Braak, 1995). The dataset spans four Brodmann (BM) brain regions: Frontal Pole (BM10), Superior Temporal Gyrus (BM22), Parahippocampal Gyrus (BM36), and Inferior Frontal Gyrus (BM44).

Data pre-processing: Genes that have a low expression on average were filtered where a cut-off of 1.00 had been estimated based on the distribution of the average expression of all genes. Then data were scaled between 0 and 1.

Pro-angiogenic gene analysis: To estimate which pro-angiogenic genes are differentially expressed in patients with a higher Braak stage; the fold change of gene expression values in patients with Braak stage 5 or higher versus the average expression values in patients with Braak stage 3 or lower for each brain region were computed independently.

Patient profiles were clustered using hierarchical clustering with ward linkage and Euclidean distance.

## Acknowledgments

We thank Prof. Stephen Payne for useful comments on the manuscript. HS is a Sir Henry Wellcome Fellow.

## Author contributions

HS and AA conceived the study. HS performed the image analysis, computational analyses, and results interpretation. AA provided the LOPAC1280 screening dataset. HS and AA thoroughly discussed and wrote the manuscript.

## Supplementary Figures

**Supplementary Figure 1.**
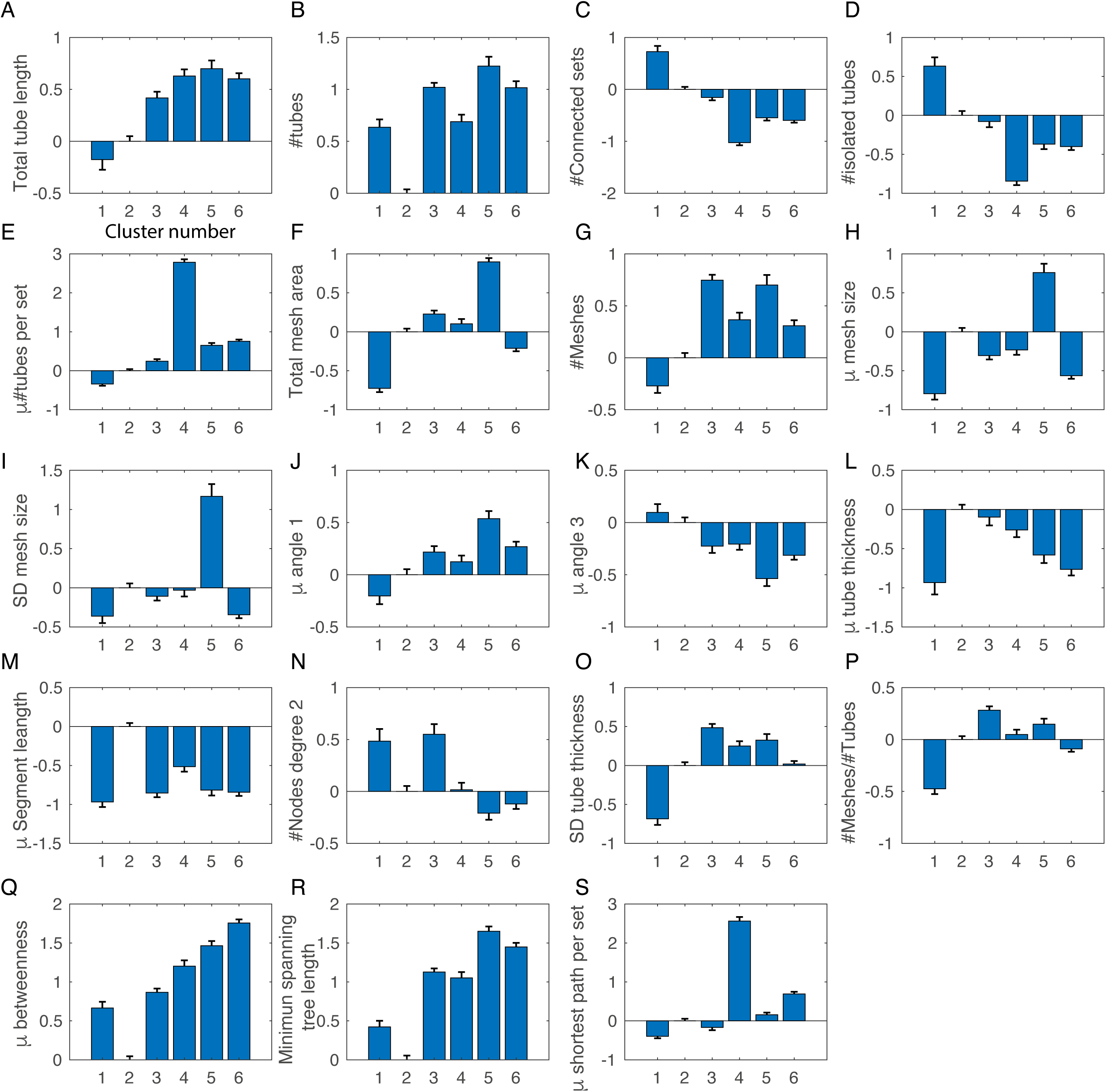
Bar charts of selected feature values for PhenoClusters 1-6.

**Supplementary Figure 2.**
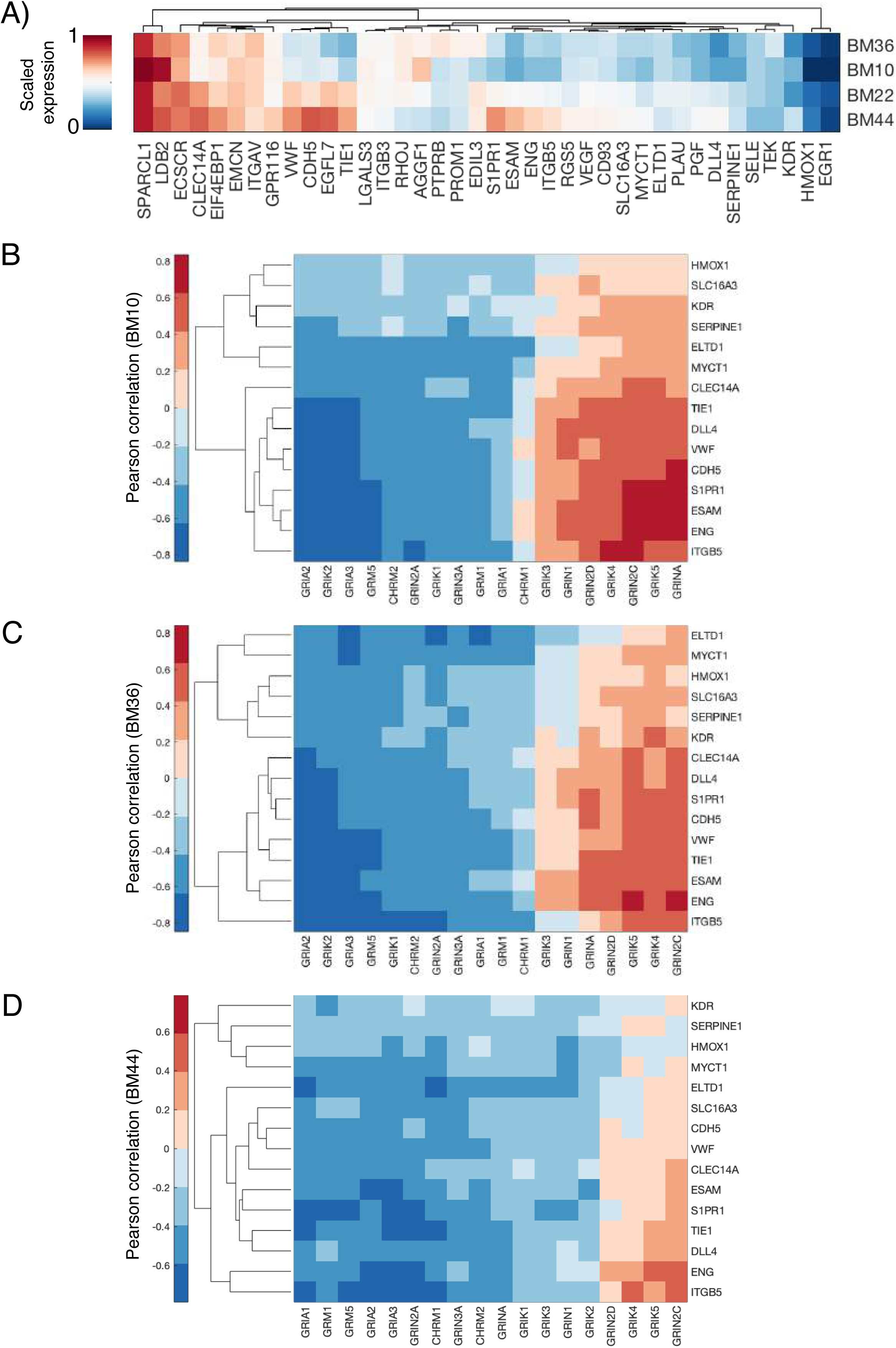
**A)** Normalised expression level of angiogenesis genes in different brain regions. **(B)** Correlation between expression levels of pro-angiogenic genes and glutamate receptor genes in brain regions BM10, BM36 and BM44.

## Supplementary tables

**Supplementary Table 1:** List of extracted endothelial network features and their categories.

**Supplementary Table 2:** Clustering results of compounds in LOPAC1280.

**Supplementary Table 3:** List of enriched mechanism of actions for each cluster and the corresponding compounds.

**Supplementary Table 4:** List of compounds that are structurally similar within clusters.

**Supplementary Table 5:** Target genes of glutamate receptors antagonists in PhenoCluster 4 and PhenoCluster 5.

